# Specialized metabolism by trichome-preferentially-expressed Rubisco and fatty acid synthase components

**DOI:** 10.1101/2022.07.12.499773

**Authors:** Wangming Ji, Sabyasachi Mandal, Yohannes H. Rezenom, Thomas D. McKnight

**Author notes:** Senior author and author for correspondence. These authors contributed equally to the paper (Wangming Ji, Sabyasachi Mandal). The author responsible for distribution of materials integral to the findings presented in this article in accordance with the policy described in the Instructions for Authors (https://academic.oup.com/plphys/pages/general-instructions) is Thomas D. McKnight.

## Abstract

Acylsugars, specialized metabolites with defense activities, are secreted by trichomes of many solanaceous plants. Several acylsugar metabolic genes (AMGs) remain unknown. We previously reported multiple candidate AMGs. Here, using multiple approaches, we characterized additional AMGs. First, we identified differentially expressed genes between high- and low-acylsugar-producing F_2_ plants derived from a cross between *Solanum lycopersicum* and *S. pennellii*, which produce acylsugars ∼1% and ∼20% of leaf dry weight, respectively. Expression levels of many known and candidate AMGs positively correlated with acylsugar amounts in F_2_ individuals. Next, we identified *lycopersicum-pennellii* putative orthologs with higher nonsynonymous to synonymous substitutions. These analyses identified four candidate genes, three of which showed enriched expression in stem trichomes compared to underlying tissues (shaved stems). Virus-induced gene silencing confirmed two candidates, *Sopen05g009610* [beta-ketoacyl-(acyl-carrier-protein) reductase; fatty acid synthase component] and *Sopen07g006810* (Rubisco small subunit), as AMGs. Phylogenetic analysis indicated that *Sopen05g009610* is distinct from specialized metabolic cytosolic reductases, but closely related to two capsaicinoid biosynthetic reductases, suggesting evolutionary relationship between acylsugar and capsaicinoid biosynthesis. Additionally, data mining revealed that orthologs of *Sopen05g009610* are preferentially expressed in trichomes of several acylsugar-producing solanaceous species. Similarly, orthologs of *Sopen07g006810* were identified as trichome-preferentially-expressed members, which form a phylogenetic clade distinct from those of mesophyll-expressed “regular” Rubisco small subunits. Furthermore, δ^13^C analyses indicated recycling of metabolic CO_2_ into acylsugars by Sopen07g006810 and shed light on how trichomes support high levels of specialized metabolite production. These findings have implications for genetic manipulation of trichome specialized metabolism in solanaceous crops, including tomato, potato, and tobacco.

## Introduction

Plant metabolites are traditionally classified into primary or central metabolites and secondary or specialized metabolites. In contrast to evolutionarily conserved primary metabolites, specialized metabolites are found in specific taxonomic groups and exhibit greater structural diversity. The building blocks of specialized metabolites are derived from products of primary metabolism; for example, alkaloids are derived from amino acids, whereas acylsugars are derived from sugar and fatty acids. The evolution of a specialized metabolic pathway requires evolution of new gene functions, which can be achieved through a variety of mechanisms, including duplication of primary metabolic genes followed by neo- or sub-functionalization and/or changes in spatio-temporal gene expression (Moghe and Last, 2015).

Specialized metabolites have important roles in plant-environment interactions. For example, acylsugars, which are nonvolatile and viscous metabolites secreted through glandular trichomes of many species in the Solanaceae (Slocombe et al., 2008; Moghe et al., 2017), provide protection against biotic and abiotic stress. These compounds contribute directly and indirectly to plant defense by providing resistance against insect herbivores (Alba et al., 2009; Leckie et al., 2016), by mediating multitrophic defense by attracting predators of herbivores through volatile short-chain aliphatic acids produced from acylsugar breakdown (Weinhold and Baldwin, 2011), and by protecting plants from microbial pathogens (Luu et al., 2017). Acylsugars also protect plants from desiccation (Fobes et al., 1985; Feng et al., 2021). These beneficial properties have led to interests in understanding acylsugar metabolism and identifying factors that control their production for breeding agronomically important crops with better resistance against insect herbivores (Bonierbale et al., 1994; Lawson et al., 1997; Leckie et al., 2012).

Acylsugars exhibit tremendous structural variation in the Solanaceae. Both branched- and straight-chain fatty acids are esterified to the sugar moiety (glucose or sucrose) to form acylsugars, and major acyl chains vary in length (C2 to C12) in different species (Kroumova et al., 2016; Moghe et al., 2017). Predominant branched-chain fatty acids include 2-methylpropanoate, 3-methylbutanoate, and 2-methylbutanoate, which are derived from branched-chain amino acids (valine, leucine, and isoleucine, respectively) (Walters and Steffens, 1990). Branched medium-chain acyl groups, such as 6-methyheptanoate and 8-methylnonanoate, are derived from branched short-chain precursors through elongation reactions mediated by either α-ketoacid (one-carbon elongation) or fatty acid synthase (FAS; two-carbon elongation) (Kroumova and Wagner, 2003). Predominant straight-chain fatty acids, such as *n*-decanoate and *n*-dodecanoate, are presumably derived from acetyl-CoA via FAS-mediated *de novo* biosynthesis (Walters and Steffens, 1990; Mandal et al., 2020). Once acyl chains are produced, specific sets of acylsugar acyltransferases (ASATs) in different species of the Solanaceae attach these aliphatic groups to different carbon positions of the sugar moiety, leading to remarkable metabolic diversity (Schilmiller et al., 2015; Fan et al., 2016; Moghe et al., 2017; Nadakuduti et al., 2017; Feng et al., 2021). Although many acylsugar metabolic genes (AMGs) have been identified in recent years, several remain unidentified and the regulation of acylsugar biosynthesis has not been well characterized, especially with regard to how solanaceous trichomes can support production of high levels of specialized metabolites.

*Solanum pennellii*, a wild relative of the cultivated tomato *Solanum lycopersicum*, is endemic to arid western slopes of the Peruvian Andes and is known for its ability to withstand extreme drought conditions. Acylsugars secreted from glandular trichomes of *S. pennellii* represent a remarkably large fraction, up to 20%, of the leaf dry weight; in contrast, *S. lycopersicum* produces only ∼1% of its leaf dry weight as acylsugars. (Fobes et al., 1985). Here, in order to identify AMGs, we first created a *S. lycopersicum* x *S. pennellii* F_2_ population and conducted transcriptomic comparisons between high- and low-acylsugar-producing F_2_ individuals to identify candidate AMGs. Next, we compared this list with our previously published list of candidate AMGs (Mandal et al., 2020). Additionally, genome-wide nonsynonymous to synonymous substitution rates (dN/dS ratio) estimation of *S. pennellii* and *S. lycopersicum* putative orthologs refined the list of candidate genes, and analysis of trichome-preferential expression identified three candidates-*Sopen05g009610*, encoding a beta-ketoacyl-(acyl-carrier-protein) reductase (SpKAR1 hereafter; a component of the FAS complex), *Sopen07g006810*, encoding a small subunit of Rubisco (SpRBCS1 hereafter), and *Sopen05g032580*, encoding an induced stolon-tip protein like member (SpSTPL hereafter). Virus-induced gene silencing (VIGS) indicated roles for both *SpKAR1* and *SpRBCS1* in acylsugar metabolism, while silencing of *SpSTPL* had no effect. The role of SpRBCS1 was further supported by δ^13^C analyses of acylsugars. Additionally, phylogenetic analyses and data mining revealed interesting evolutionary aspects which suggested that orthologs of SpKAR1 and SpRBCS1 are involved in specialized metabolism in other plants.

## Results

### Transcriptomic comparison between high- and low-acylsugar-producing F_2_ individuals of *S. lycopersicum* x *S. pennellii*

*S. pennellii* accession LA0716 produces copious amount of acylsugars (∼20% of leaf dry weight), whereas *S. lycopersicum* cv. VF36 accumulates considerably lower amount (∼1% of leaf dry weight) (Fobes et al., 1985). The interspecific F_1_ hybrid (LA4135) accumulates low levels of acylsugars (<3% of leaf dry weight). To identify candidate AMGs, we first analyzed acylsugar accumulation in an F_2_ population derived from the cross between VF36 and LA0716. Of the 114 F_2_ plants, 24 accumulated acylsugars >10% of their leaf dry weight, whereas 62 accumulated acylsugars <3% of their leaf dry weight (Figure 1A). Next, using RNA-seq, we identified genes that were differentially expressed between 10 high- and 10 low-acylsugar-producing F_2_ individuals (hereafter referred to as HIGH-F_2_ and LOW-F_2_, respectively). A total of 20,160 *S. pennellii* genes were selected after minimum-expression-level filtering, and 331 differentially expressed genes (DEGs) were identified; of these, 134 and 197 DEGs showed higher and lower expression levels, respectively, in the HIGH-F_2_ group compared to the LOW-F_2_ group (Supplemental Data Set 1). Enrichment analysis indicated that gene ontology (GO) terms such as “acyltransferase activity” (GO:0016747), “fatty acid metabolic process” (GO:0006631), and “active transmembrane transporter activity” (GO:0022804) were over-represented in the list of 331 DEGs (Supplemental Figure S1).

**Figure 1.**
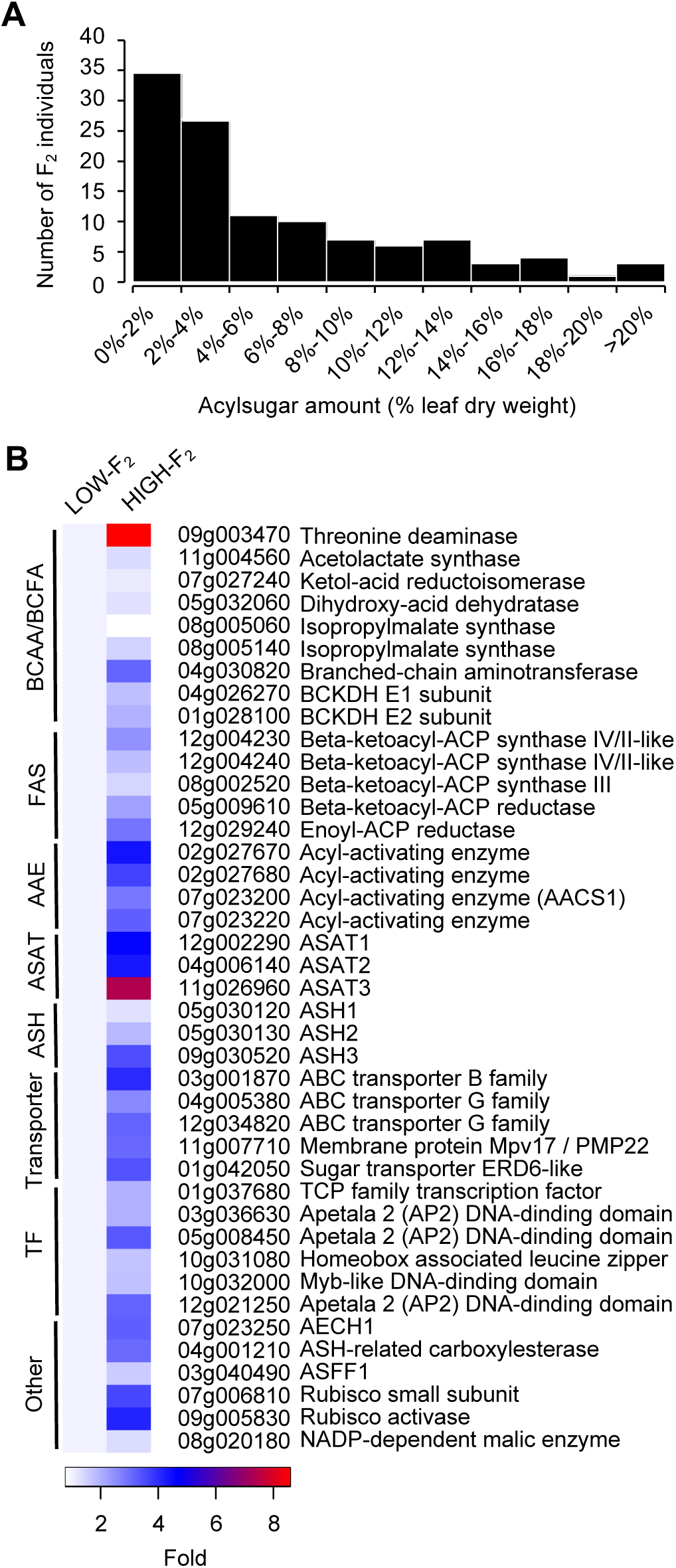
Acylsugar accumulation and expression of known and candidate acylsugar metabolic genes (AMGs) in a *Solanum lycopersicum* VF36 x *S. pennellii* LA0716 F_2_ population. A, Histogram showing acylsugar amount distribution among 114 F_2_ plants. For each plant, three replicates were used to measure acylsugar amount. B, Heatmap showing relative expression levels (set to one-fold in the LOW-F_2_ group) of genes with known and putative roles in acylsugar metabolism. *S. pennellii* gene identifier numbers (Sopen IDs) are given with annotations. BCAA= branched-chain amino acid; BCFA= branched-chain fatty acid; FAS= fatty acid synthase component; AAE= acyl-activating enzyme; ASAT= acylsugar acyltransferase; ASH= acylsugar acylhydrolase; TF= transcription factor; BCKDH= branched-chain keto acid dehydrogenase; AACS1= acylsugar acyl-CoA synthetase 1; AECH1= acylsugar enoyl-CoA hydratase 1; ASFF1= acylsucrose fructofuranosidase 1.

Of the 331 DEGs, 73 were also differentially expressed between high- and low-acylsugar-producing accessions of *S. pennellii* (Mandal et al., 2020). Many genes with known and putative roles in acylsugar metabolism showed higher expression levels in the HIGH-F_2_ group compared to the LOW-F_2_ group (Figure 1B), which validated our RNA-seq approach. These genes encode branched-chain fatty acid metabolic proteins, FAS components, acyl-activating enzymes (also known as acyl-CoA synthetases), a mitochondrial/peroxisomal membrane protein (Sopen11g007710), three ATP-binding cassette transporters, three ASATs, transcription factors, and a Rubisco small subunit. Additionally, known and putative flavonoid metabolic genes, such as *Sopen11g003320* (UDP-glucose:catechin glucosyltransferase) and three sequential genes on chromosome 6 (*Sopen06g034810, Sopen06g034820*, and *Sopen06g034830*; myricetin *O*-methyltransferase; (Kim et al., 2014)), which are strongly co-expressed with AMGs (Mandal et al., 2020), were also found in the list of DEGs (Supplemental Data Set 1).

### Genome-wide dN/dS ratio estimation of *S. pennellii* and *S. lycopersicum* putative orthologs and refinement of candidate AMG list

In addition to difference in gene expression levels, difference in protein-coding sequence may also contribute to difference in acylsugar accumulation capabilities between *S. pennellii* and *S. lycopersicum*. Compared to primary metabolic genes, specialized metabolic genes evolve faster and exhibit higher ratio of nonsynonymous to synonymous substitution rates (dN/dS ratio) (Moore et al., 2019). Therefore, as an additional approach for identifying candidate AMGs, we performed a genome-wide dN/dS ratio analysis (Yang and Nielsen, 2000) of *S. pennellii* and *S. lycopersicum* putative orthologs to identify genes that are under positive selection. Using reciprocal BLAST, 19,984 putative ortholog pairs were selected (Supplemental Data Set 2), and the yn00 maximum-likelihood method (Yang, 1997) yielded a genome-wide mean dN/dS ratio of 0.3273 (Figure 2A). A total of 732 genes with dN/dS >1.0 were considered to be under positive selection (Supplemental Data Set 3).

**Figure 2.**
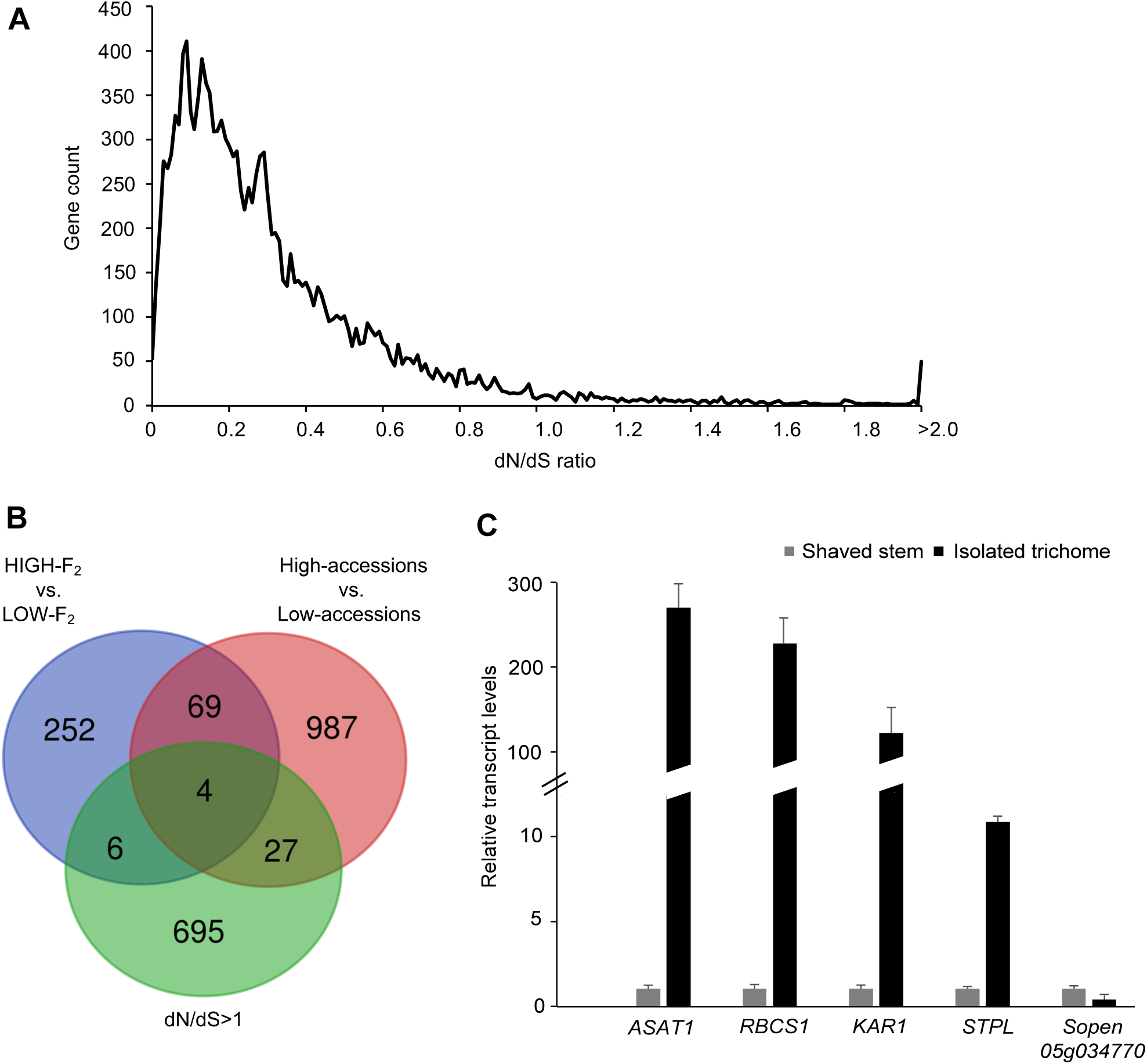
Selection of candidate AMGs. A, Distribution of nonsynonymous to synonymous substitution rate ratios (dN/dS ratios) of putative ortholog pairs from *S. pennellii* and *S. lycopersicum*. B, Venn diagram showing the intersections of three sets of candidate AMGs: (1) differentially expressed genes (DEGs) between high- and low-acylsugar-producing *S. pennellii* accessions (Mandal et al., 2020; red circle), (2) DEGs between high- and low-acylsugar-producing F_2_ plants (HIGH-F_2_ and LOW-F_2_, respectively; purple circle), and (3) genes with dN/dS > 1 between *S. pennellii* and *S. lycopersicum* putative orthologs (green circle). Four genes were identified at the intersections of these three sets. C, Relative expression levels of the four candidate AMGs [*SpRBCS1* (*Sopen07g006810*), *SpKAR1* (*Sopen05g009610*), *SpSTPL* (*Sopen05g032580*) and *Sopen05g034770*] in isolated stem trichomes and underlying tissues of shaved stems (normalized to one-fold) in *S. pennellii* LA0716. *SpASAT1* (*Sopen12g002290*), the ortholog of *S. lycopersicum* trichome tip-cell-expressed *ASAT1* (Fan et al., 2016), was included for comparison. Error bars indicate SE (*n* = 5 individual plants).

To narrow our focus further, we next compared lists of candidate AMGs obtained from three approaches: (A) 331 DEGs identified in our analysis of the F_2_ population, (B) 1,087 DEGs we previously reported from transcriptomic analysis between high- and low-acylsugar-producing *S. pennellii* accessions (Mandal et al., 2020), and (C) 732 genes with dN/dS >1.0. Four genes (*Sopen07g006810, Sopen05g009610, Sopen05g032580*, and *Sopen05g034770*) occurred in all three sets (Figure 2B, Table 1). Next, we measured trichome-enriched expression of these four candidates, since many AMGs are expressed in trichome tip-cells (Ning et al., 2015; Schilmiller et al., 2015; Fan et al., 2016; Fan et al., 2020). Transcript levels of *Sopen07g006810* (*SpRBCS1*), *Sopen05g009610* (*SpKAR1*), and *Sopen05g032580* (*SpSTPL*) were 220-, 110- and 13-fold, respectively, higher in isolated stem trichomes than in underlying tissues of shaved stems in *S. pennellii* accession LA0716. On the other hand, expression of *Sopen05g034770* was lower in trichomes than in shaved stems (Figure 2C). Because of this, and the fact that expression of *Sopen05g034770* is inversely correlated with acylsugar amount (Table 1), this gene was not analyzed further.

**Table 1.** Four genes identified at the intersections of three sets of candidate AMGs

| Gene ID | High vs. Low accessions log<sub>2</sub>FC | HIGH-F<sub>2</sub> vs. LOW-F<sub>2</sub> log<sub>2</sub>FC | dN/dS value | dN/dS rank | Annotation |
| --- | --- | --- | --- | --- | --- |
| Sopen07g006810 | 5.01 | 1.85 | 2.91 | 29 | Rubisco small subunit |
| Sopen05g009610 | 3.32 | 1.15 | 2.8 | 35 | Beta-ketoacyl-(acyl-carrier-protein) reductase |
| Sopen05g032580 | 2.97 | 2.1 | 1.52 | 227 | Induced stolon tip protein-like |
| Sopen05g034770 | -3.48 | -1.89 | 1.59 | 199 | Uncharacterized protein |

Log<sub>2</sub>FC indicates log<sub>2</sub> (fold-change). Positive and negative log<sub>2</sub>FC values indicate higher and lower expression levels, respectively, in high-acylsugar-producing *S. pennellii* accessions (Mandal et al., 2020) or HIGH-F<sub>2</sub> group. dN/dS rank represents the corresponding rank of selected gene in 732 putative orthologs with dN/dS > 1.

### *In vivo* functional validation of candidate AMGs

To determine whether *SpRBCS1, SpKAR1* and *SpSTPL* are involved in acylsugar biosynthesis, these three candidate genes were targeted in *S. pennellii* LA0716 for VIGS using tobacco rattle virus (TRV)-based silencing vectors (Dong et al., 2007). VIGS resulted in significant downregulation of target genes (82%, 54%, and 94% reduction in transcript levels for *SpRBCS1, SpKAR1*, and *SpSTPL*, respectively; Supplemental Figure S2, A and B), and leaf surface metabolites were analyzed by liquid chromatography-mass spectrometry (LC-MS). Compared to a group of control plants (empty TRV vectors), total acylsugar levels decreased by 23% (*P* < 0.05) in VIGS-*SpRBCS1* plants (Figure 3, A and B; Supplemental Data Set 4). In contrast, we did not observe any statistically significant changes in total acylsugar amount upon suppression of *SpKAR1* or *SpSTPL* (Figure 3A).

**Figure 3.**
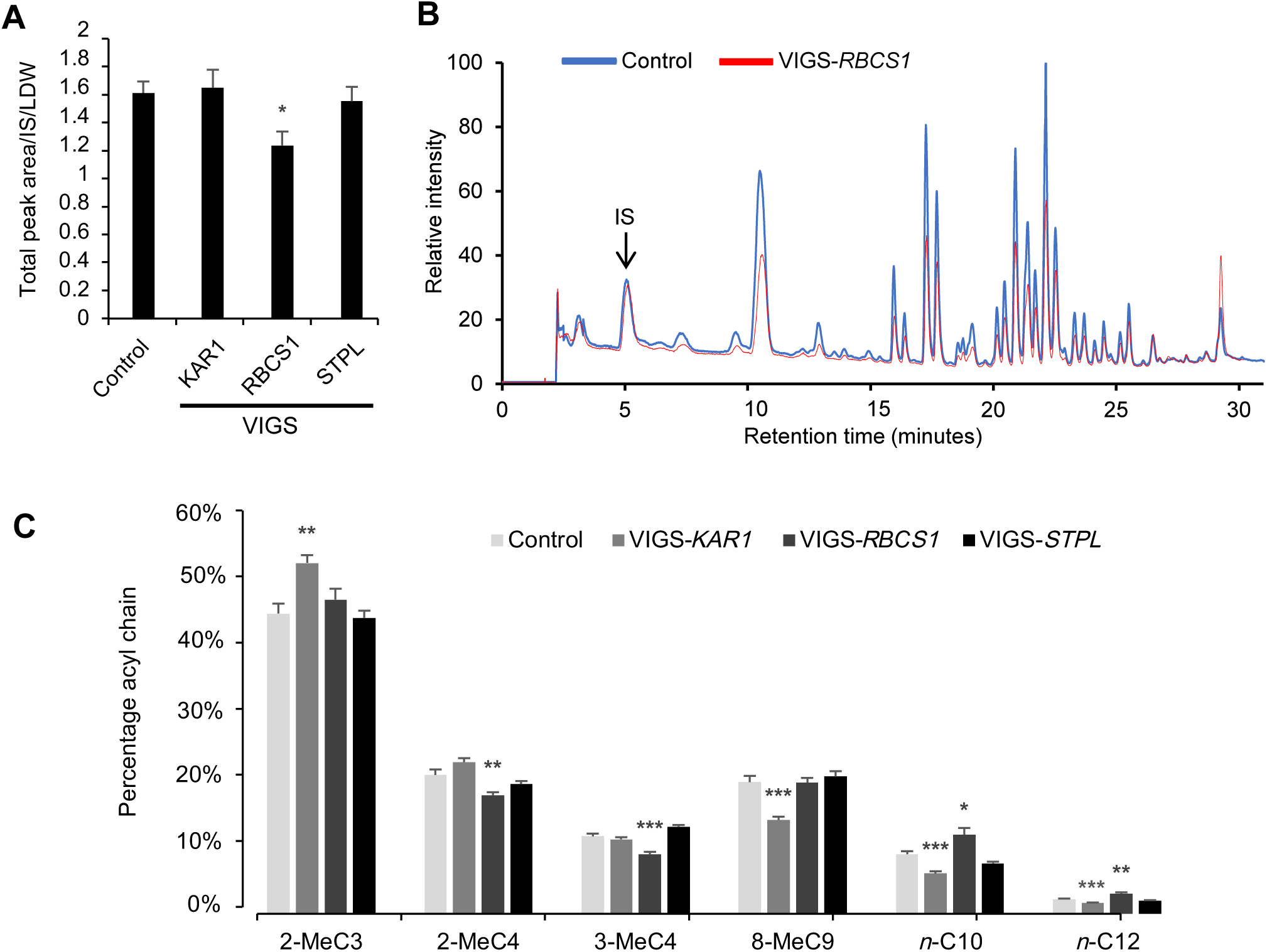
Virus-induced gene silencing (VIGS) of three trichome-preferentially-expressed candidate AMGs in *S. pennellii* LA0716. A, Acylsugar quantification by liquid chromatography– mass spectrometry (LC-MS). To quantify acylsugar amounts, chromatogram peak areas were normalized by internal standard (IS) area and leaf dry weight (LDW). Error bars indicate SE (*n* = 10, 12, 11, and 12 individual plants for control, VIGS-*KAR1*, VIGS-*RBCS1* and VIGS-*STPL* groups, respectively; * *P* < 0.05; Dunnett’s test). B, Representative chromatograms (normalized by IS area and LDW) showing acylsugar peaks in control and VIGS-*RBCS1* plants. Acylsugar peaks are listed in Supplemental Data Set 4. C, Acylsugar acyl chain composition analysis by gas chromatography-mass spectrometry (GC-MS). Predominant acyl chains are shown. Me= methyl; C3-C12 indicate acyl chain length (for example, 2-MeC3 and *n*-C10 indicate 2-methlpropanoate and *n*-decanoate, respectively). Error bars indicate SE (*n* = 10, 12, 11, and 12 individual plants for control, VIGS-*KAR1*, VIGS-*RBCS1* and VIGS-*STPL* groups, respectively; * *P* < 0.05, ** *P* < 0.01, *** *P* < 0.001; Dunnett’s test).

Despite the lack of influence on total acylsugar amount, silencing of *SpKAR1* led to significant reductions in acylsugar medium-chain fatty acids (18%, 23%, and 20% reductions for 8-methylnonanoate, *n*-decanoate, and *n*-dodecanoate, respectively), which is consistent with its predicted role in medium-chain fatty acid biosynthesis (Figure 3C). Silencing of *SpRBCS1* resulted in 20% reductions in C5 acyl chains (2- and 3-methylbutanoate). No noticeable morphological differences were observed between control and silenced plants (Supplemental Figure S2C), which indicates that these effects on acylsugar phenotypes were not indirect consequences of abnormal plant growth and development caused by VIGS. Transcript enrichment in trichomes and VIGS results together indicate roles of SpRBCS1 and SpKAR1 in acylsugar metabolism. On the other hand, no statistically significant changes in acyl chain profile were observed upon silencing of *SpSTPL*.

### Phylogenetic analysis of SpKAR1

Short-chain dehydrogenases/reductases (SDRs) constitute a large protein superfamily of NAD(P)(H)-dependent oxidoreductases; members exhibit low levels of sequence identity, but they share a Rossmann-fold motif for nucleotide binding (Kavanagh et al., 2008). Many SDRs are involved in biosynthesis of specialized metabolites, such as sesquiterpene zerumbone in *Zingiber zerumbet* (Okamoto et al., 2011), diterpene momilactone in *Oryza sativa* (Shimura et al., 2007), monoterpenoids in glandular trichomes of *Artemisia annua* (Polichuk et al., 2010), monoterpenoid constituents of essential oils in peppermint and spearmint (Ringer et al., 2005), phenolic monoterpenes in the Lamiaceae (Krause et al., 2021), and steroidal glycoalkaloids and saponins in *Solanum* species (Sonawane et al., 2018). SpKAR1 (322-aa) belongs to the SDR family and shares several common motifs, such as the N-terminal cofactor binding motif TGxxxGxG, a downstream structural motif (C/N)NAG, the active site motif YxxxK, and the catalytic tetrad N-S-Y-K, with members of the SDR “classical” subfamily (Kavanagh et al., 2008) (Supplemental Figure S3). However, phylogenetic analysis suggested that SpKAR1 is closer to *Escherichia* and *Synechocystis* FabG [beta-ketoacyl-(acyl-carrier-protein) reductase] than other specialized metabolic SDRs mentioned earlier (Figure 4A; Supplemental Figure S4). Additionally, these plant SDRs are predicted to have a cytosolic location, whereas SpKAR1 is predicted to be localized to the chloroplast [TargetP (https://services.healthtech.dtu.dk/service.php?TargetP-2.0), WoLF PSORT (https://www.genscript.com/wolf-psort.html), and MultiLoc2 (https://abi-services.informatik.uni-tuebingen.de/multiloc2/webloc.cgi)]. Furthermore, cytochrome P450 monooxygenase (CYP) activities are closely associated with these specialized metabolic SDRs, whereas SpKAR1 activity presumably is not associated with CYP products. These results together indicate that SpKAR1 is a component of the FAS complex in the chloroplast, and it is phylogenetically distant from other specialized metabolic SDRs mentioned earlier.

**Figure 4.**
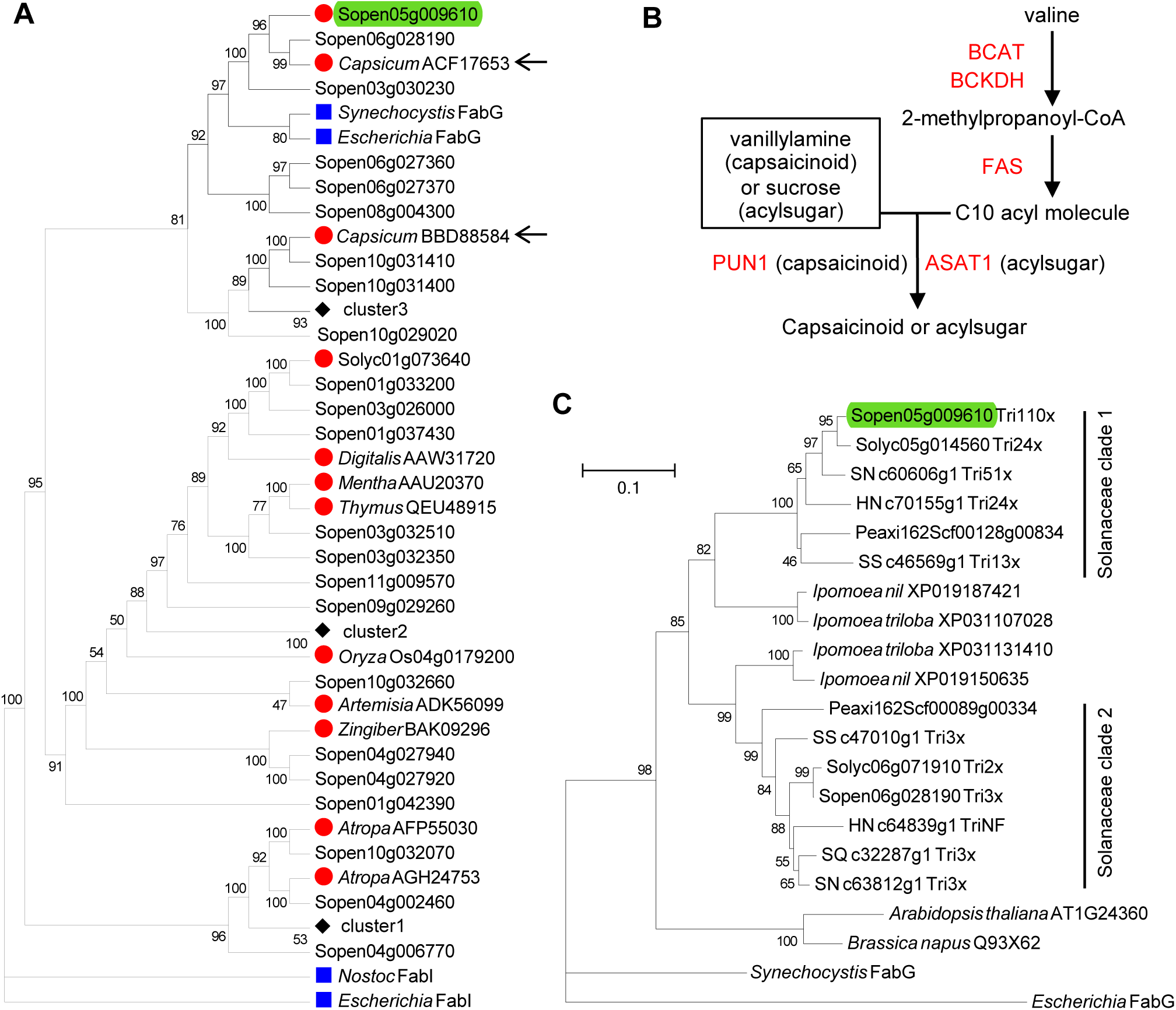
Phylogenetic analyses of SpKAR1. A, Maximum-likelihood tree (topology) of SpKAR1 (Sopen05g009610; highlighted) and related short-chain dehydrogenases/reductases (SDRs). Red circles indicate plant specialized metabolic SDRs. Blue squares indicate bacterial sequences. “Sopen” numbers indicate sequences from *Solanum pennellii*. Sequences from other species are given with GenBank accession numbers. Black diamonds indicate more than one “Sopen” sequences, which were clustered to save space; complete tree is given in Supplemental Figure S4. Two sequences related to capsaicinoid biosynthesis are indicated by arrows. Bootstrap values from 1000 replicates are shown on the nodes. B, Similarity between capsaicinoid and acylsugar biosynthetic pathways. Metabolism of leucine, isoleucine, and straight-chain fatty acids in acylsugar pathway are not shown here. Single arrows do not necessarily indicate single enzymatic steps. Enzymes are in red font. BCAT= branched-chain aminotransferase; BCKDH= branched-chain keto acid dehydrogenase; FAS= fatty acid synthase; PUN1= pungent gene 1; ASAT1= acylsugar acyltransferase 1. C, Neighbor-joining tree of SpKAR1 (highlighted) and its homologs in the Solanaceae. Sequences from four non-solanaceous plant species [*Ipomoea triloba* and *I. nil* (Convolvulaceae); *Arabidopsis thaliana* and *Brassica napus* (Brassicaceae)] and two bacteria (*Synechocystis* and *Escherichia*) are also included. Bootstrap values from 1000 replicates are shown. Tree is drawn to scale, with branch lengths measured in the number of substitutions per site. Tri110x indicates 110-fold higher expression in isolated trichomes compared to underlying tissues (NF= not found in HN_c64839g1). RT-qPCR was used for “Sopen” sequences. Trichome-enriched expression data (based on RNA-seq) for sequences in five other species were obtained from Ning et al., 2015 (Solyc= *S. lycopersicum*) and Moghe et al., 2017 (SN= *S. nigrum*; SQ= *S. quitoense*; HN= *Hyoscyamus niger*; SS= *Salpiglossis sinuata*). Peaxi= *Petunia axillaris*. Sopen03g030230 (138-aa) and its putative orthologs were not included because they have long deletions and insertions. Maximum-likelihood tree is given in Supplemental Figure S5.

*Capsicum annuum* is a solanaceous species that produces capsaicinoid specialized metabolites, but not acylsugars. Two SDRs from *Capsicum annuum* showed close phylogenetic relationships with SpKAR1, and we investigated the similarity between capsaicinoid and acylsugar biosynthetic pathways. In both pathways, valine is converted to 2-methylpropanoyl-CoA, which is then elongated to C10 acyl molecules via FAS-mediated reactions (Figure 4B). This suggests a common evolutionary origin for acyl chain elongation steps in both acylsugar and capsaicinoid biosynthesis.

Ning et al. (2015) reported trichome-enriched expression data for *S. lycopersicum* genes, and Moghe et al. (2017) reported similar data in four additional acylsugar-producing solanaceous species-*S. nigrum, S. quitoense, Hyoscyamus niger*, and *Salpiglossis sinuata*. We mined these publicly available datasets and investigated if *SpKAR1* orthologs exhibit trichome-enriched expression. Phylogenetic analysis identified two distinct clades, and members of one clade are preferentially expressed in trichomes (Figure 4C; Supplemental Figures S5 and S6). This suggests that SpKAR1 orthologs have a role in trichome acylsugar biosynthesis in other solanaceous species.

### Phylogenetic analysis of SpRBCS1

Rubisco is composed of eight large subunits (encoded by a single *RBCL* gene on the plastid genome) and eight small subunits (encoded by a multigene family *RBCS* on the nuclear genome). Among the five annotated *RBCS* genes in *S. pennellii* (three on chromosome 2 and one on each of chromosome 3 and 7), only *Sopen07g006810* (*SpRBCS1*) exhibited noticeable trichome-enriched expression based on reverse transcription-quantitative PCR (Supplemental Figure S7A). In shaved stems, *SpRBCS1* had extremely low, if any, level of expression (if *SpRBCS1* is trichome-specific, residual expression could be due to incomplete shaving of stem trichomes). On the other hand, other *RBCS* members showed high expression levels in shaved stems (∼1000-fold higher than *SpRBCS1*). Additionally, protein sequence analysis revealed that other RBCS members, which share >90% sequence identity with each other, share low level of homology (<55% identity) with SpRBCS1 (Supplemental Figure S7, A and B). Furthermore, *Sopen07g006810*-*Solyc07g017950* ortholog pair had dN/dS ratio of 2.91, whereas ortholog pairs of other RBCS members showed dN/dS ratios less than 0.32 (Supplemental Data Set 3), which is expected from highly conserved, mesophyll-expressed “regular” *RBCS* members (presumably involved in primary metabolism). These results indicated that SpRBCS1 is distinct from other RBCS members. This was supported by phylogenetic analysis, which in conjunction with data mining, showed that orthologs of *SpRBCS1* are preferentially expressed in trichomes of solanaceous members (average 112-fold in six species where trichome-enriched expression data are available; Figure 5; Supplemental Figures S8 and S9). Interestingly, *SpRBCS1* was placed in a monophyletic clade with *Nicotiana tabacum RBCS-T*, which is expressed in trichome tip cells (Laterre et al., 2017). Orthologs of SpRBCS1 were found outside the Solanaceae (including *Oryza sativa*), but not in Arabidopsis, which lacks glandular trichomes.

**Figure 5.**
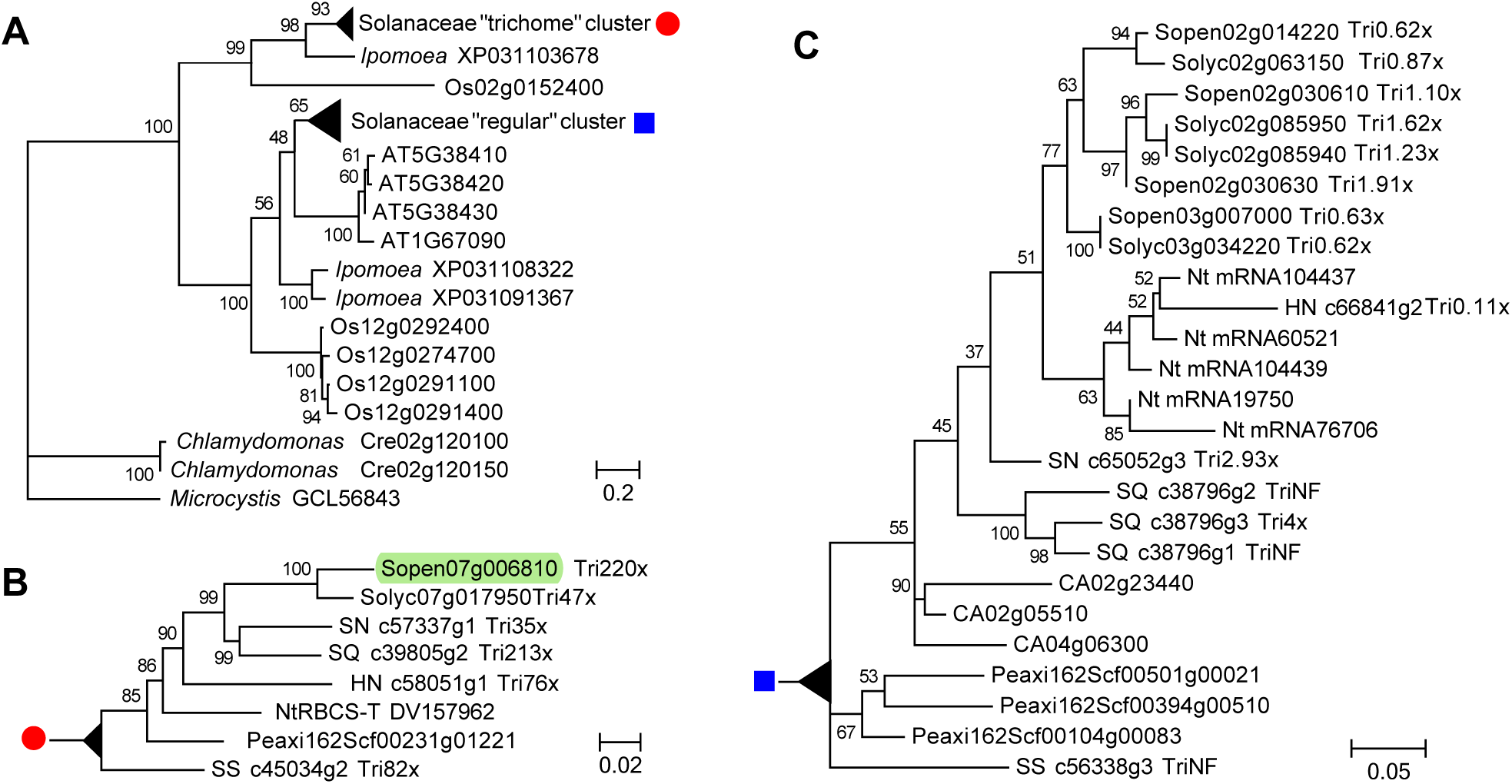
Phylogenetic analysis of SpRBCS1 (Sopen07g006810; highlighted). A, Maximum-likelihood tree of Rubisco small subunits. Solanaceous sequences were combined into two clusters (indicated by a red circle and a blue square) to save space. Neighbor-joining tree is given in Supplemental Figure S8. Bootstrap values from 1000 replicates are shown on the nodes. Tree is drawn to scale, with branch lengths measured in the number of substitutions per site. Os= *Oryza sativa*; AT= *Arabidopsis thaliana*. GenBank accession numbers are indicated for *Ipomoea triloba* and *Microcystis aeruginosa*. B and C, Expanded Solanaceae “trichome” cluster (B) and expanded Solanaceae “regular” cluster (C). Tri220x indicates 220-fold higher expression in isolated trichomes compared to underlying tissues (NF= not found). RT-qPCR was used for *Solanum pennellii* (Sopen) sequences. Trichome-enriched expression data (based on RNA-seq) for sequences in five other species were obtained from Ning et al., 2015 (Solyc= *S. lycopersicum*) and Moghe et al., 2017 (SN= *S. nigrum*; SQ= *S. quitoense*; HN= *Hyoscyamus niger*; SS= *Salpiglossis sinuata*). Peaxi= *Petunia axillaris*; CA= *Capsicum annuum*; Nt= *Nicotiana tabacum*.

### δ^13^C analyses of acylsugars

Based on a previous metabolomics study (Balcke et al., 2017), it was hypothesized that Rubisco in trichomes mainly recycles CO_2_ released by the high metabolic rate in these cells. In *S. pennellii* trichomes, where production of acylsugars is high, three steps in this pathway release CO_2_: acetolactate synthase, isopropylmalate dehydrogenase, and branched-chain ketoacid dehydrogenase. If trichome Rubisco is responsible for recycling CO_2_, then acylsugars (and other metabolites in trichomes) will contain some carbon that has undergone at least two rounds of fixation, first in the bulk of the plant to provide primary metabolites, and again in the trichome to recover CO_2_ released during production of branched chain fatty acids. One consequence of this re-fixation would be that acylsugars contain even less ^13^C than the rest of the plant due to isotopic fractionation at each fixation (the lighter ^12^C isotope would be favored because of a lower activation energy). To test this hypothesis, we measured fractionation of carbon isotopes (δ^13^C; reported in parts per thousand ‰) in both secreted acylsugars and plant tissues presumably without acylsugars (shaved stems). Acylsugars and shaved stems showed δ^13^C of - 33.15‰ and -28.99‰, respectively (Figure 6A). The difference in δ^13^C between these two sample types indicates that acylsugars contain carbons that have undergone additional rounds of fixation compared to non-acylsugar metabolites. Additionally, δ^13^C values from *S. pennellii* samples (acylsugars and shaved stems) are comparable to δ^13^C value from another member of the Solanaceae that was reported previously (-30.7‰ in *Nicotiana tabacum*) (Smith and Epstein, 1971).

**Figure 6.**
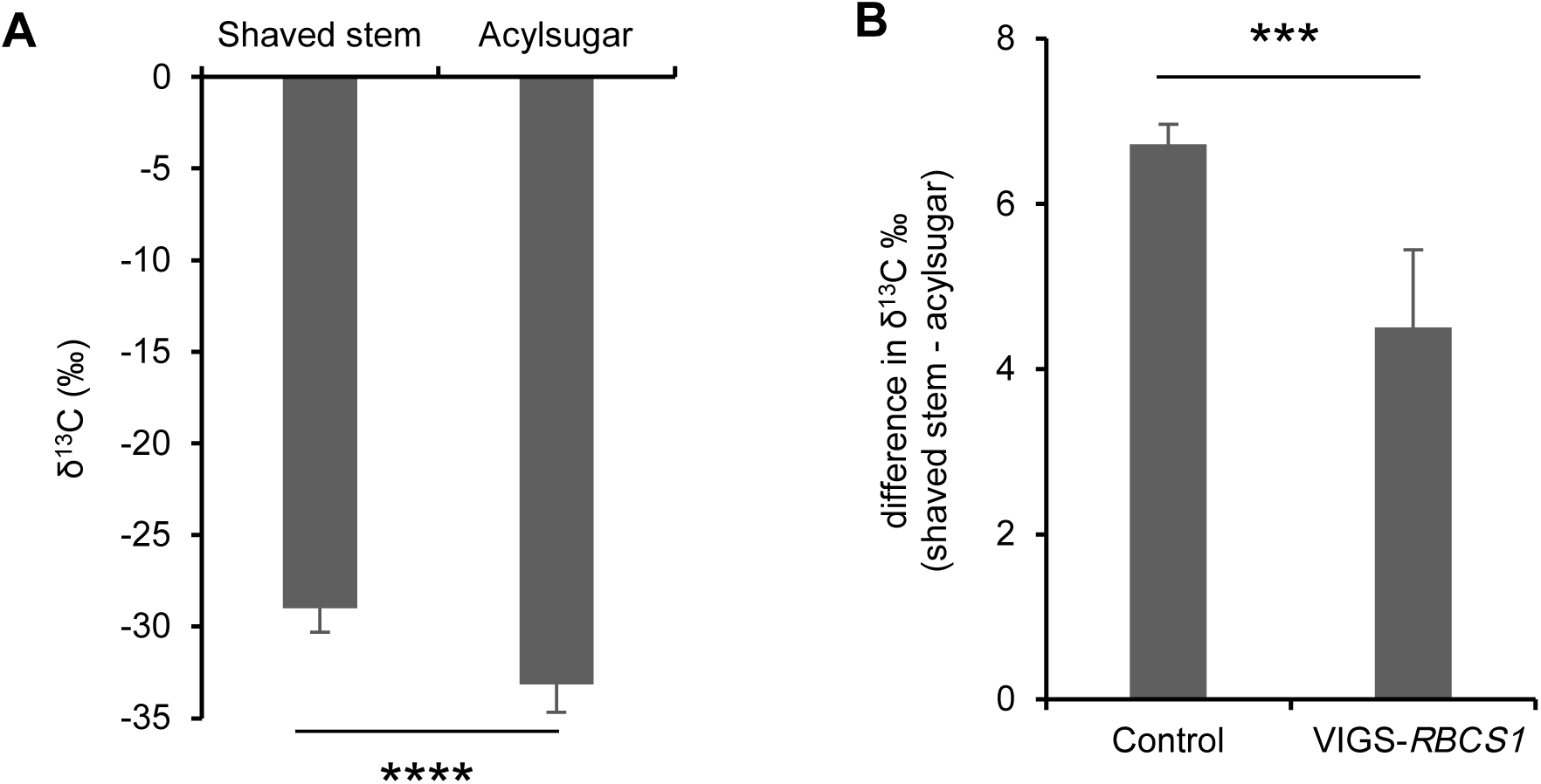
δ^13^C analyses. A, Difference in δ^13^C values between shaved stems and secreted acylsugars. Error bars indicate SE (*n* = 10 individual plants; **** *P* < 0.0001; Welch *t*-test). B, VIGS of *SpRBCS1* reduces the difference in δ^13^C values between shaved stems and acylsugars. Error bars indicate SE (*n* = 5 and 10 individual plants for control and VIGS-*RBCS1* groups, respectively; *** *P* < 0.001; Welch *t*-test).

Next, we hypothesized that the trichome-preferentially-expressed *SpRBCS1* is responsible for this re-fixation of carbons into acylsugars. To test our hypothesis, we determined the difference in δ^13^C between shaved stems and acylsugars (δδ^13^C) for two groups of plants-control and VIGS-*SpRBCS1*. Compared to a control group, VIGS led to a reduction in δδ^13^C (Figure 6B), which confirmed the role of SpRBCS1 in supplying re-fixed carbons for acylsugar biosynthesis.

## Discussion

Acylsugars are secreted by glandular trichomes of several plant families, including Martyniaceae (Asai et al., 2010), Caryophyllaceae (Asai et al., 2012), Geraniaceae (Sakai et al., 2013), and Solanaceae, where they have been best studied. A detailed knowledge about genes involved in regulating acylsugar amount and acyl chain profile (Ben-Mahmoud et al., 2018) is required for successful crop breeding programs and metabolic engineering of acylsugar production. Here, we report two trichome-preferentially-expressed genes involved in acylsugar biosynthesis.

### Multiple approaches to identify and validate candidate AMGs

Many enzymes involved in acylsugar biosynthesis have been identified in recent years (Ning et al., 2015; Schilmiller et al., 2015; Fan et al., 2016; Schilmiller et al., 2016; Moghe et al., 2017; Nadakuduti et al., 2017; Fan et al., 2020; Mandal et al., 2020; Feng et al., 2021; Lou et al., 2021). To identify additional enzymes in this pathway, and hopefully regulatory proteins, we used a multipronged strategy to find genes whose expression and evolutionary patterns suggested they could be involved. To refine our previously reported list of candidate AMGs (Mandal et al., 2020), we looked for genes that were differentially expressed between high- and low-acylsugar-producing F_2_ individuals derived from *S. lycopersicum* x *S. pennellii*, and also genes that appeared to be rapidly evolving as determined by the dN/dS ratio. The intersection of three sets of candidate AMGs yielded four candidates (Figure 2B). One candidate, *Sopen05g034770*, had lower expression in high-acylsugar-producing F_2_ individuals than in low-acylsugar-producing F_2_ individuals and also lower expression in trichomes relative to underlying tissue (Table 1; Figure 2C). Both of these patterns are opposite to what we would expect for genes involved in acylsugar biosynthesis. Therefore, this gene was not studied further, but the hypothetical protein encoded by this gene may act as a negative regulator of the pathway. Putative roles of the remaining three trichome-preferentially-expressed candidates in acylsugar biosynthesis were tested using VIGS, which confirmed *SpKAR1* and *SpRBCS1* as AMGs (Figure 3). Phylogenetic analyses and data mining revealed that orthologs of both *SpKAR1* (Figure 4C) and *SpRBCS1* (Figure 5) are trichome-preferentially-expressed members in the Solanaceae, suggesting roles for these orthologs in acylsugar biosynthesis. Additionally, *SpKAR1* and *SpRBCS1* exemplify duplication of highly-conserved primary metabolic genes followed by spatial regulation of gene expression as a driver of evolution of acylsugar metabolism.

### Function and phylogeny of SpKAR1

In plants, *de novo* fatty acid biosynthesis takes place predominantly in plastids, which use a multicomponent type II FAS system that catalyzes the extension of the growing acyl chain. The plastidic FAS has four well-characterized enzymatic components: a beta-ketoacyl-(acyl-carrier-protein) reductase (KAR), a beta-hydroxyacyl-(acyl-carrier-protein) dehydrase, an enoyl-(acyl-carrier-protein) reductase, and three isozymes of beta-ketoacyl-(acyl-carrier-protein) synthases (KAS I, II, and III). KAR catalyzes one of the steps of the core four-reaction cycle of the FAS-mediated chain elongation process. Close phylogenetic relationship with bacterial FabG (KAR) and its predicted chloroplast location indicate that SpKAR1 is a component of the plastid FAS, and it is distinct from other specialized metabolic cytosolic SDRs (Figure 4A). VIGS showed that SpKAR1 is required for acylsugar straight-chain fatty acid (SCFA) biosynthesis (Figure 3C), and we previously reported two trichome-preferentially-expressed KAS genes that are involved in SCFA biosynthesis (Mandal et al., 2020). These results corroborate the role of FAS in acylsugar SCFA biosynthesis.

Elongation of acetyl-CoA to SCFAs by FAS complex presumably takes place in plastids; however, it is not clear if elongation of 2-methylpropanoyl-CoA (isobutyryl-CoA; produced in mitochondria from valine) to 8-methylnonanoyl-CoA (one of the major branched-chain acyl groups) occurs in mitochondria or chloroplast (Figure 4B). For both capsaicinoid (Mazourek et al., 2009) and acylsugar (Slocombe et al., 2008) biosynthesis, it has been proposed that 2-methylpropanoyl-CoA is transported from mitochondria to chloroplast, where it is elongated by the plastid FAS complex. Recently, a dual-localized KAR (AT1G24360) was reported in Arabidopsis (Guan et al., 2020), and AT1G24360 is phylogenetically closely related to SpKAR1 (Figure 4C; Supplemental Figure S5). This suggests the possibility that plastidic-SpKAR1 is involved in the biosynthesis of acylsugar SCFAs, whereas mitochondrial-SpKAR1 is involved in the elongation of 2-methylpropanoyl-CoA; this would allow 2-methylpropanoyl-CoA to be elongated without being transported to plastid.

### Capsaicinoid and acylsugar biosynthetic pathways

In *Capsicum*, 8-methyl-6-nonenoyl-CoA (C10) and 8-methylnonanoyl-CoA (C10) are attached to an aromatic compound derived from phenylalanine to generate capsaicin and dihydrocapsaicin, respecetively, which are two of the most potent capsaicinoids (Mazourek et al., 2009). These C10 acyl molecules are presumably derived from 2-methylpropanoyl-CoA via FAS-mediated reactions, and this segment of the capsaicinoid biosynthetic pathway is similar to the acylsugar biosynthetic pathway (Figure 4B). Attachment of acyl groups to an aromatic compound (in case of capsaicinoid biosynthesis) or sucrose (in case of acylsugar biosynthesis) is catalyzed by BAHD superfamily transferases: PUN1 in capsaicinoid biosynthesis (Stewart et al., 2005) and ASATs in acylsugar biosynthesis (Schilmiller et al., 2015; Fan et al., 2016). Moghe et al. (2017) reported that PUN1 and ASATs share a common evolutionary ancestor, and our phylogenetic analysis revealed that SpKAR1 is closely related to two reductases involved in capsaicinoid biosynthesis (FAS-mediated acyl chain elongation steps) (Figure 4A). These results indicate close phylogenetic relationships between segments of capsaicinoid and acylsugar biosynthetic pathways. Additionally, acyl chain length is an important factor in determining both pungency of capsaicinoids and potential in defense activity of acylsugars (Ben-Mahmoud et al., 2018). Taken together, these findings will be useful in future metabolic engineering of capsaicinoid and acylsugar production.

### Function and phylogeny of SpRBCS1

Phylogenetic analysis clustered *SpRBCS1* with other solanaceous trichome-preferentially-expressed *RBCS* members (Figure 5), including *Nicotiana tabacum* tip-cell-expressed *RBCS-T* (Laterre et al., 2017), and also with rice *OsRBCS1*, which is expressed in several tissues other than leaf blade (major photosynthetic organ) (Morita et al., 2014). Additionally, based on enzymatic properties, both RBCS-T and OsRBCS1 were found to be distinct from “regular” RBCS members in respective species. These findings indicate specialized functions for SpRBCS1 and its orthologs in special cell/tissue types.

Trichome metabolites can accumulate at noticeably high levels; for example, acylsugars in *S. pennellii* and duvatrienediol in *N. tabacum* can accumulate up to 20% and 15%, respectively, of leaf dry weight (Fobes et al., 1985; Severson et al., 1985). This indicates high metabolic activities and CO_2_ release in trichome cells (for example, enzymatic steps catalyzed by acetolactate synthase, isopropylmalate dehydrogenase, and branched-chain ketoacid dehydrogenase complex generate CO_2_ during acylsugar production). However, due to thick cell walls and cuticle, it has been suggested that trichomes have limited gaseous exchange with the outside and fix little atmospheric CO_2_ (Balcke et al., 2017). In order to support continued high metabolic activities, trichomes may require re-fixation of metabolic CO_2_ with a Rubisco that is active in high CO_2_ and low pH conditions. Biochemical assays indicate that *N. tabacum* RBCS-T has been adapted to such conditions (Laterre et al., 2017). These results encouraged us to investigate, using δ^13^C analysis, if SpRBCS1 is involved in re-fixation of metabolic CO_2_ into acylsugars.

Fractionation of carbon isotopes during photosynthesis occurs predominantly during the carboxylation reaction (carbon fixation) catalyzed by Rubisco, and it leads to a preferential enrichment of one stable isotope over another. Photosynthates contain less of the ^13^C than the ^12^C due to kinetic isotope effects-the lighter ^12^C isotope is preferentially incorporated into products because it has a higher energy state (a lower activation energy) (Smith and Epstein, 1971). We used this information about carbon isotope fractionation to analyze metabolites, and tested the hypothesis based on Balcke et al. (2017) that in trichomes, Rubisco mostly re-fixes metabolic CO_2_, which are incorporated into specialized metabolites (predominantly acylsugars in case of *S. pennellii*). Our δ^13^C examination of acylsugars and shaved stems showed recycling of metabolic CO_2_ into acylsugars (Figure 6A). Additionally, VIGS confirmed that the trichome-preferentially-expressed *SpRBCS1* is responsible for this re-fixation (Figure 6B). Tomato trichomes receive carbon mostly from leaf sucrose (Balcke et al., 2017); however, re-fixation of metabolic CO_2_ would allow trichomes to maintain a pH homeostasis and also to improve carbon utilization for sustained high level production of specialized metabolites.

## Materials and methods

### Plant materials and growth conditions

Seeds of *Solanum pennellii* LA0716 and the F_1_ hybrid (LA4135) of *S. lycopersicum* VF36 x *S. pennellii* LA0716 were obtained from the C.M. Rick Tomato Genetics Resource Center (University of California, Davis). The LA4135 was self-pollinated for generating the F_2_ population. Seeds were treated with 1.2% (w/v) sodium hypochlorite for 20 minutes and rinsed with deionized water three times before placing on moist filter paper in petri dishes. After germination, seedlings were transferred to soil and grown in a growth chamber (16-hour photoperiod; 24°C/20°C day/night temperature; 150 µMol m^-2^ s^-1^ photosynthetically active radiation; 75% relative humidity).

### Acylsugar collection from F_2_ individuals

Secreted acylsugars were collected from three young leaves of 10-week-old F_2_ individuals as three replicates by dipping them in ethanol for 2-3 seconds. Ethanol was completely removed by evaporation until dryness in a fume hood. Acylsugar amount was determined as a proportion of leaf dry weight. Leaf dry weights were measured after drying in a 70°C oven for one week.

### RNA sequencing (RNA-seq)

10 high-acylsugar-producing F_2_ individuals (acylsugar amounts >14% of leaf dry weight) and 10 low-acylsugar-producing F_2_ individuals (acylsugar amounts <1% of leaf dry weight) were used in the HIGH-F_2_ versus LOW-F_2_ transcriptome comparison. After removing surface metabolites with ethanol for 2-3 seconds, young leaves were immediately frozen with liquid nitrogen, and stored at -80°C until further use. Total RNA was isolated from leaves using the RNAqueous Total RNA Isolation Kit (Thermo Fisher Scientific), and the genomic DNA was removed using the TURBO DNA-Free Kit (Thermo Fisher Scientific). RNA-seq libraries of polyA^+^-selected samples were prepared using TruSeq Stranded mRNA Library Preparation Kit LT (Illumina). After quality control, libraries were sequenced on the HiSeq 4000 (Illumina) 150×150-bp paired-end sequencing platform according to the manufacturer’s specifications at the Texas A&M Genomics and Bioinformatics Service Center, College Station.

### Differential gene expression analysis

Approximately 31 to 48 million (average 36 million) paired-end reads were generated from RNA-seq libraries. Sequencing reads were processed using Trimmomatic v0.32 (Bolger et al., 2014) with the following settings: ILLUMINACLIP:TruSeq3-PE-2.fa:2:30:10, LEADING = 20, TRAILING = 20, SLIDINGWINDOW = 4:20, MINLEN = 100. Approximately 68% of the reads in each library passed the trimming filter. The trimmed reads were then mapped to the *S. pennellii* LA0716 genome v2.0 (Bolger et al., 2014) using TopHat2 v2.1.0 (Kim et al., 2013) with the following parameters: -library-type = fr-firststrand, -mate-inner-dist = 0, -mate-std-dev = 50, -read-realign-edit-dist = 1000, -read-edit-dist = 2, -read-mismatches = 2, -min-anchor-len = 8, -splice-mismatches = 0, -min-intron-length = 50, -max-intron-length = 50,000, -max-insertion-length = 3, -max-deletion-length = 3, -max-multihits = 20, -min-segment-intron = 50, - max-segment-intron = 50,000, -segment-mismatches = 2, -segment-length = 25. 70-85% of the trimmed reads were mapped to the *S. pennellii* genome. Aligned reads from TopHat2 were counted for each gene using HTseq package version 0.6.1 (Anders et al., 2015) with the following parameters: -f bam, -r name, -s reverse, -m union, -a 20. The count files were used to identify differentially expressed genes (DEGs) using edgeR version 3.32.1 (Robinson et al., 2010). Fragments per kilobase per million mapped reads (FPKM) value for each gene in each sample was called with rpkm command in edgeR program. Genes with more than one count per million (CPM) in at least two samples were used for differential gene expression analysis. DEGs were identified when *P*, corrected for multiple testing, was less than 0.05 (false discovery rate < 0.05), and fold change was greater than 2.

### Identification of putative orthologs and dN/dS estimation

To identify putative orthologs, we performed an all-versus-all reciprocal BLAST between annotated genes of *S. pennellii* v2.0 (Bolger et al., 2014) and *S. lycopersicum* ITAG2.3 (Tomato Genome, 2012) with the following settings: minimum percentage identity = 70, minimum percentage query coverage = 50. Putative orthologs were aligned with ClustalW, and the alignment information was converted into codon alignments using PAL2NAL (Suyama et al., 2006). Genome-wide dN/dS ratios were calculated between putative ortholog pairs using the yn00 maximum likelihood method in the PAML package (Yang, 1997; Yang and Nielsen, 2000).

### Determination of trichome-enriched expression

Reverse transcription-quantitative PCR (RT-qPCR) was used to measure trichome-enriched expression of selected genes in *S. pennellii* LA0716, as described in Mandal et al. (2020). RT-qPCR primers are given in Supplemental Table S1.

### Virus-induced gene silencing (VIGS)

VIGS was performed using the tobacco rattle virus (TRV)-based vectors (Dong et al., 2007) in *S. pennellii* LA0716. VIGS constructs were designed using the Solanaceae Genomics Network VIGS tool (http://vigs.solgenomics.net/) and were cloned into the pTRV2-LIC vector, in the antisense orientation, to target selected genes. *Agrobacterium tumefaciens* strain GV3101 harboring pTRV1, pTRV2 constructs, and empty pTRV2 were grown overnight with 50 mg/mL kanamycin and 10 mg/mL gentamicin at 28°C. Cultures were centrifuged at 8,000g for 5 min at 4°C, and cells were washed and resuspended in infiltration buffer (10 mM MES pH 5.5, 10 mM MgCl_2_, and 200 μM of acetosyringone). Cell suspensions were incubated at room temperature for 3 hours, and different pTRV2 cultures were mixed with equal volumes of pTRV1 cultures to reach final OD_600_ = 1 before infiltration at the first true leaf stage with a needleless syringe. Plants were grown in a chamber with conditions mentioned earlier for approximately six weeks. Silencing of target genes were assessed with RT-qPCR using primers that were designed outside the VIGS-targeted regions. VIGS primers are listed in Supplemental Table S1.

### Chromatography–mass spectrometry analysis

Secreted acylsugars from control and VIGS plants were quantified with liquid chromatography-mass spectrometry (LC–MS). Acylsugars on leaf surface were collected from similar-sized young leaves by submerging them in 10 ml of extraction solvent [acetonitrile:isopropanol:water (3:3:2, v/v/v) with 0.1% formic acid; 100 μM propyl 4-hydroxybenzoate was used as the internal standard], followed by gentle mixing for 2 minutes. Extracted samples were analyzed using Q Exactive Focus coupled with Ultimate 3000 RS LC unit (Thermo Fisher Scientific) and Exactive Series 2.8 SP1/Xcalibur 4.0 software. Acylsugars were separated by injecting 10 µL of sample into Acclaim 120 (2.1 × 150 mm; 3 µm) C18 column (Thermo Fisher Scientific) that was housed at 30°C. 0.1% formic acid was used as eluent A and acetonitrile with 0.1% formic acid was used as eluent B in the mobile phase. Flow rate was set at 300 µL/min with the following gradient: 0– 3 minutes, 40% B; 3-23 minutes, 40-100% B; 23-28 minutes, hold 100% B; 28.1-31 minutes, hold 40% B. The Q Exactive Focus HESI source was operated in full MS in negative electrospray ionization (ESI) mode. Other parameters were set as follows: sheath and auxiliary gas flow rates-35 and 10 arbitrary units, respectively; spray voltage-3.3 kV; S-Lens RF level-50 v. The transfer capillary temperature and the auxiliary gas heater temperature were held at 320°C and 350°C, respectively. Parallel reaction monitoring (PRM) mode was used for targeted MS/MS of acylsugars. Relative abundances of acylsugars were determined by dividing total peak areas of all detected acylsugars with peak area of the internal standard and leaf dry weight. Dry weights of the extracted leaves were measured after one week of drying in a 70°C oven.

Acylsugar acyl chain profiles were analyzed with gas chromatography-mass spectrometry (GC–MS) after performing transesterification reaction, as described in Mandal et al. (2020).

### Phylogenetic analysis

Sequences were obtained from the Solanaceae Genomics Network, GenBank, and the NCBI website. For *S. nigrum, S. quitoense, Hyoscyamus niger*, and *Salpiglossis sinuata, de novo* assembled transcriptomes (Moghe et al., 2017) were used to collect sequences, which were translated in six possible frames (https://web.expasy.org/translate/) to obtain protein sequences with longest open reading frames. MAFFT (Katoh and Standley, 2013) was used for multiple sequence alignment with BLOSUM62 matrix, gap extend penalty value of 0.123, and gap opening penalty value of 1.53. ModelFinder (Kalyaanamoorthy et al., 2017) was used to compare substitution models, and the best model of protein evolution was selected based on lowest Bayesian Information Criterion scores (LG+I+G4 for Figure 4A, JTT+G4 for Supplemental Figure S5, and LG+G4 for Figure 5). IQ-TREE 2 (Minh et al., 2020) was used to construct maximum-likelihood based phylogenetic trees. MEGA X (Kumar et al., 2018) was used to construct neighbor-joining phylogenetic trees applying Jones-Taylor-Thornton (JTT) model. Uniform rates among sites were used, and pairwise deletion was used for gaps/missing data. Bootstrap values were obtained from 1000 replicates.

### Stable isotope analysis

δ^13^C analysis was performed at the Texas A&M Stable Isotope Geosciences Facility using methods described in McDermott et al. (2019).

### Accession numbers

Sequence data from this article can be found in the GenBank data libraries under accession numbers ON920880 (SpRBCS1; Sopen07g006810), ON920881 (SpRBCS2; Sopen02g014220), ON920882 (SpRBCS3; Sopen02g030610), ON920883 (SpRBCS4; Sopen02g030630), ON920884 (SpRBCS5; Sopen03g007000), ON920885 (SpKAR1; Sopen05g009610), ON920886 (SpKAR2; Sopen06g028190), ON920887 (SpSTPL; Sopen05g032580), and ON920888 (hypothetical protein; Sopen05g034770). RNA-seq reads used in this study were submitted to the NCBI Sequence Read Archive under the accession number PRJNA818092 (BioProject ID).

## Supplemental data

**Supplemental Figure S1**. Enrichment analysis of gene ontology (GO) terms associated with 331 differentially expressed genes (DEGs) between high- and low-acylsugar-producing F_2_ individuals.

**Supplemental Figure S2**. Virus-induced gene silencing (VIGS) of candidate genes in *Solanum pennellii* LA0716.

**Supplemental Figure S3**. Multiple sequence alignment of SpKAR1 (Sopen05g009610) and its homologs, including specialized metabolic short-chain dehydrogenases/reductases (SDRs).

**Supplemental Figure S4**. Maximum-likelihood phylogenetic tree of SpKAR1 (Sopen05g009610) and related short-chain dehydrogenases/reductases (SDRs).

**Supplemental Figure S5**. Maximum-likelihood phylogenetic tree of SpKAR1 (Sopen05g009610) and its homologs in the Solanaceae.

**Supplemental Figure S6**. Multiple sequence alignment of SpKAR1 (Sopen05g009610) and its homologs in the Solanaceae.

**Supplemental Figure S7**. Analyses of RBCS members in *Solanum pennellii*.

**Supplemental Figure S8**. Neighbor-joining tree of SpRBCS1 (Sopen07g006810) and its homologs.

**Supplemental Figure S9**. Multiple sequence alignment of SpRBCS1 (Sopen07g006810) and its homologs.

**Supplemental Table S1**. List of primers used in this study.

**Supplemental Data Set 1**. Differentially expressed genes (DEGs) between HIGH-F_2_ and LOW-F_2_ groups.

**Supplemental Data Set 2**. Reciprocal Best Hits (RBH) between *Solanum pennellii* and *S. lycopersicum* annotated sequences.

**Supplemental Data Set 3**. 732 putative ortholog pairs between *Solanum pennellii* and *S. lycopersicum* with dN/dS > 1.

**Supplemental Data Set 4**. Acylsugars from *Solanum pennellii*.

## Acknowledgments

We sincerely thank Charlie Johnson and the staff of the Texas A&M Genomics and Bioinformatics Center for performing Illumina sequencing, Michael Dickens and the staff of the Texas A&M High Performance Research Computing Platform for providing computational resources and assistance, and Christopher Maupin of the Texas A&M Stable Isotope Geosciences Facility for assistance with δ^13^C analysis.

## Author contributions

WJ and SM designed and performed experiments, analyzed data, and wrote the manuscript. YHR designed and performed chromatography–mass spectrometry analyses, and reviewed the manuscript. TDM designed experiments, analyzed data, and wrote the manuscript. All authors read and approved the final manuscript.

